# Distinguishing between interaction and dispersion effects in the analysis of gene-environment interaction

**DOI:** 10.1101/2020.09.08.287888

**Authors:** Benjamin W. Domingue, Klint Kanopka, Travis T. Mallard, Sam Trejo, Elliot M. Tucker-Drob

**Affiliations:** Graduate School of Education, Stanford University & Center for Population Health Sciences, Stanford Medicine; Graduate School of Education, Stanford University; Department of Psychology, University of Texas at Austin; La Follette School of Public Affairs & Department of Sociology, University of Wisconsin–Madison; Department of Psychology & Population Research Center, University of Texas at Austin

**Keywords:** gene-by-environment interaction, gene × environment interaction, GxE, G×E, vQTL, heteroscedasticity

## Abstract

Genotype-by-environment interaction (GxE) occurs when the size of a genetic effect varies systematically across levels of the environment and when the size of an environmental effect varies systematically across levels of the genotype. However, total variance in the phenotype may shift as a function of the moderator irrespective of its etiology such that the *proportional* effect of the predictor is constant. We expand the traditional GxE regression model to directly account for heteroscedasticity associated with both the genotype and the measured environment. We then derive a test statistic, *ξ*, for inferring whether GxE can be attributed to an effect of the moderator on the dispersion of the phenotype. We apply this method to identify genotype-by-birth year interactions for Body Mass Index (BMI) that are distinguishable from general secular increases in the variance of BMI or associations of the genetic predictors (both PGS and individual loci) with BMI variance. We provide software for analyzing such models.

## 1 Introduction

Genotype-by-Environment interaction (GxE) studies test whether the genotype-phenotype association varies in magnitude across the range of a measured environmental variable and whether the environment-phenotype association varies in magnitude across the range of the measured genotype. Investigations of GxE have been of particular interest in the study of complex traits, leading to a variety of methods for estimating polygenic GxE [1, 2]. Development of these analytic approaches coincides with a large increase in the availability of genomic data, particularly in the context of population-based studies that also contain a broad range of phenotypes and environmental measures. Here, we consider circumstances where either the genetic or the environmental measure selected for GxE is associated with the magnitude of variance in the target phenotype. Such heteroscedasticity introduces a key interpretational problem to results from standard GxE approaches: any observed GxE may arise as an artifact of the fact that the total variance in the phenotype shifts across the range of the genetic or environmental moderator, irrespective of etiology. We resolve this problem by expanding the standard regression-based model for testing GxE in both single locus and polygenic score settings by introducing a simple yet flexible approach to directly model heteroscedastic residuals, and we develop a test statistic for inferring whether GxE can be plausibly attributed to a more general effect of the moderator on the dispersion of the phenotype.

All else equal, when the dispersion of a phenotype is directly controlled by the level of the moderator, the standard regression-based model will identify a significant interaction between that moderator and *any* correlate of the phenotype, whether genetic or non-genetic. For example, when the effectiveness, intensity, or dosage of a phenotype-altering intervention varies proportionally to baseline levels of the phenotype (e.g. when hours of individualized education are assigned based on previous year’s test scores; or when intensity of a weight-loss intervention is determined from baseline body mass index), then the variance of the phenotype is expected to increase in response to the intervention, and the unstandardized effect sizes for all correlates of baseline levels of the phenotype are expected to increase. Similarly, when a genetic variant confers greater plasticity in a phenotype, then the variance of the phenotype is expected to differ across alleles (such that the variant is identified as a variance quantitative trait locus, vQTL). We would then expect the variant to moderate the effect sizes for all correlates of that phenotype proportionally to one another. Only when a moderator acts preferentially on specific mechanisms of variation in the phenotype are interactions with various predictors expected to depart from expectations that follow from differences in total variance across the range of the moderator.

Figure 1 illustrates how heteroscedasticity across the range of a measured environmental moderator (E; left panel) or genetic moderator (G; right panel) can produce the impression of GxE. In the left panel, we illustrate a scenario in which a PGS is associated with a constant proportion of variance in the phenotype across the range of the measured environment. That the proportion of variance explained by the PGS at any given location on the x-axis is constant indicates that the expected percentile location within the distribution of the phenotype will also be constant across the range of *E* (assuming that the shape, but not dispersion, of the distribution remains constant across the range of *E*). Nevertheless, a clear fan-spreading effect is observed, incorrectly implying that the PGS is increasingly penetrant at higher levels of E. The right panel represents an analogous scenario with respect to a single genetic variant that moderates the variance of the phenotype (i.e a vQTL [3–7]). In this scenario, the environment is associated with a constant proportion of variance in the phenotype across the range of the genotype, but appears to become increasingly predictive of the phenotype as the effect allele count increases as a result of heteroscedasticity. By directly modeling this heteroscedasticity, we are able to distinguish between scenarios such as these—in which GxE is an epiphenomenon of variance modulation—from those representing more meaningful patterns of GxE.

**Figure 1:**
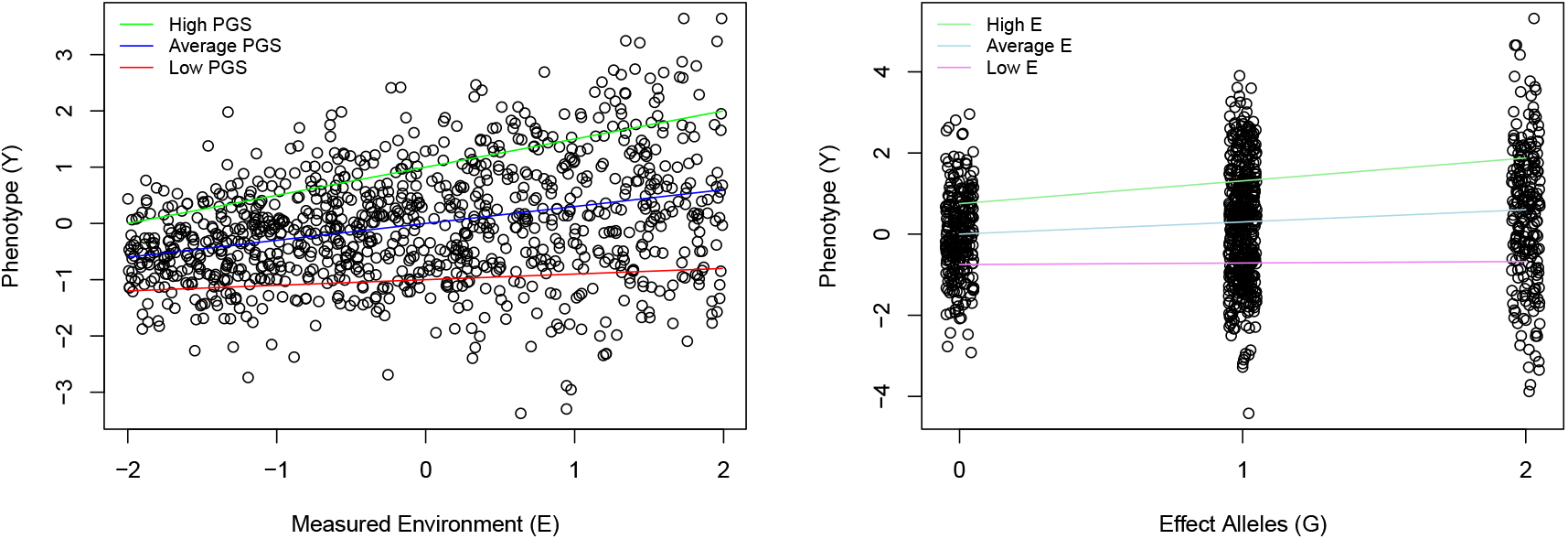
Hypothetical examples of an Environmental moderator (left) or a genetic moderator (right) of total variance in a phenotype producing the impression of GxE. Left: Residuals of the regression of the Phenotype on the Measured Environment are heteroscedastic. high, average, and low polygenic scores (PGSs) are associated with constant percentiles of the phenotype at any given location along the x axis. However, because variance of the phenotype expands across the range of the Measured Environment, the PGS accounts for increasing unstandardized variance across this range. Right: A variance quantitative trait locus (vQTL) in which the effect allele is associated with greater variance in the phenotype. Unstandardized scores on the phenotype that are associated with high, average, and low levels of the measured environment (E) become more distinct with increasing number of effect alleles. However, because the total variance in the phenotype expands across the x axis, the *percentile* locations of these scores within each genotype (0, 1, or 2) is constant across all levels of the genotype.

We develop and validate a flexible heteroscedastic regression model for testing GxE. We use this model to elucidate the well-studied interaction between birth cohort and genetics linked to body mass index (BMI). We test whether both PGS-by-Birth Year and SNP-by-Birth Year interactions on BMI can be attributed to more general increases in the total dispersion of BMI over historical time. That is, we ask whether birth year is associated specifically with amplification of genetic risk for BMI or more generally with amplification of all variation in BMI.

## 2 Methods

In settings wherein focus is on a measured genotype, as indexed by a single-locus allele count or a polygenic score (PGS), the standard regression model for GxE is

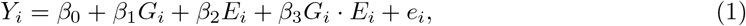

where *Y_i_* is the phenotype for person *i*, *G_i_* is the measured genotype, *E_i_* is the measured environment, and *e_i_* is an error term. Under this model, the effect of *G_i_* on *Y_i_* is allowed to vary as a linear function of *E_i_*, as indexed by the regression coefficient *β*_3_. However, a crucial assumption is that the residuals, *e_i_*, are homoscedastic across all levels of *E_i_* (e.g., 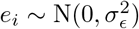).

The standard regression model given by Eqn 1 can be expanded so as to relax the assumption of homoscedasticity of residuals across *E_i_* by specifying the following *environmental heteroscedasticity model*,

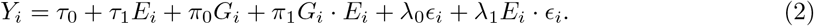

Here, *τ*_1_ is the main effect of the measured environment, *π*_0_ is the main effect of *G_i_*, *π*_1_ is the GxE analogue to *β*_3_ in Eqn 1, λ_0_ is the main effect of the error *ϵ_i_* (which we assume, without loss of generality given λ_0_, to have unit variance), and λ_1_ is the coefficient used to index heteroscedasticity. Rather than simply adjusting standard errors and p values of the standard regression model (Eqn 1) using a robust estimation method, this model explicitly estimates heteroscedasticity as a model parameter. As we articulate next, directly modelling heteroscedasticity allows us to overcome specification bias that would result from estimating a standard homoscedastic GxE model, even with a robust estimator. We are purposeful in choosing a function form for the heteroscedasticity term, *E_i_* · *ϵ_i_*, that directly parallels that specified for the GxE term, *G_i_ · E_i_*.

The heteroscedasticity model is a flexible model allowing for both conventional GxE and heteroscedasticity. In order to test whether a more general pattern of heteroscedasticity as a function of *E_i_* is sufficient to account for the observed GxE, we propose a restricted form version of this model that does not directly model GxE but instead allows for the total variance of the phenotype to vary as a function of *E_i_*. This *scaling model* represents a scenario wherein GxE—in the sense of a significant estimate of *β*_3_ in Eqn 1—arises as a function of differences in the total variance of *Y_i_* as a function of *E_i_*. We choose this nomenclature given that the model emphasizes the importance of the *scale* of the outcome. Under this model, the proportional contribution of *G_i_* is constant over the range of *E_i_*, but the scale of *Y_i_* systematically varies across the range of *E_i_*; i.e., *E_i_* acts as a “dimmer” [8].

The scaling model for moderation of the variance of *Y_i_* as a function of *E_i_* takes the form

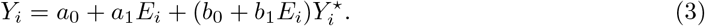

Here, 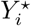 is an unobserved factor representing unexplained variation in *Y_i_* incremental of *E_i_*. The *b*_0_ + *b*_1_*E_i_* coefficient on the *Y*^⋆^ term produces heteroscedasticity in *Y_i_* as a function of *E_i_*. We let

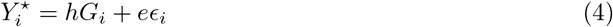

where *G* is a measured genotype standardized to have unit variance and *ϵ_i_* is an unobserved error term (we assume it is also scaled to have unit variance). Finally, we identify the units of *Y*^⋆^ by specifying 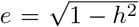. The penetrance of the measured genotype is thus controlled via the relative magnitudes of *h* and *e*. Note, in particular, that in terms of 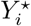, the penetrance of the measured genotype is constant. The test that we introduce relies on this property; i.e., when properly scaled, the penetrance of the measured genotype to *Y_i_* is constant. Suppose that *b*_1_ = 0. In that case, *Y_i_* is affected solely by environmental and genetic main effects. If *b*_1_ ≠ 0, then the role of 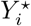 with respect to *Y_i_* varies as a function of *E*, and as we elucidate next, so do the raw (but not relative) contributions of *G_i_* and *ϵ_i_* to *Y*.

In Section A of the supplemental information (SI), we show that the environmental heteroscedasticity model reduces to the environmental scaling model when

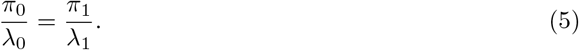

We use this observation to derive the test statistic *ξ_E_* and associated hypothesis test of whether an empirical estimate of GxE is distinguishable from the scaling model as

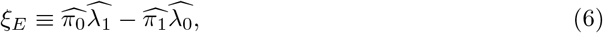

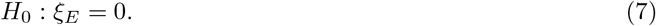

We provide parallel heteroscedasticity and scaling models for moderation of the variance of *Y_i_* as a function of *G_i_* (leading to an analogous test statistic, *ξ_G_*) in Section A.3 of SI.

We conduct a variety of simulation studies (see Section C of SI). In summary, results of these simulation studies suggest that *ξ_E_* is a reliable indicator of whether the scaling model is the basis for GxE. When a test of the statistic fails to reject *H*_0_ : *ξ_E_* =0 and there is significant GxE (*π*_1_) observed, we cannot rule out that the scaling model is driving observed GxE. When the test suggests rejection of *H*_0_ : *ξ_E_* =0, alternative forms of GxE are implicated. Software to estimate the models considered here is described in Section D of SI.

## 3 Heteroscedasticity in context of BMI-linked genetics and birth year

We apply our heteroscedastic GxE model to the well-studied interaction between birth year and genetics linked to body mass index (BMI) [9–12], a literal textbook example (see Section 11.3 of [13]). We test whether previous reports of increasing penetrance of PGS for BMI for later-born birth cohorts can be plausibly attributed to more general increases in the total dispersion of BMI, including variation unique of its genetic etiology, across birth cohorts [14, 15]. We thus ask whether birth year is associated specifically with amplification of genetic risk for BMI or more more generally with amplification of all variation in BMI. We then go on to apply our heteroscedastic GxE model to examine individual locus-by-year interactions for BMI. To illustrate how our approach can be used to conduct SNP-level analyses, we apply our technique using the top hits from a recent BMI GWAS [16] to investigate whether they are associated with heteroscedastic GxE.

### 3.1 Polygenic Score Analysis

We consider GxE in the context of a long-running cohort study—the Health and Retirement Study (HRS) [17]—of US adults over 50 and their spouses (N=11,586). Respondents are interviewed every two years and data are collected on their early-life, anthropometrics, socioeconomic status, and mental and physical well-being. Previous examinations of increases in the penetrance of the BMI polygenic score across birth year have used this data [9–11]. We use a polygenic score for BMI [16] as constructed by the HRS [18]; further description of the data are in E of SI.

Results are presented in Table 2. We observe considerable evidence for stronger prediction of BMI by the PGS for more recently born individuals (i.e., *π*_1_ > 0); this is consistent with earlier findings indicating secular increases in the penetrance of a BMI PGS [9–11]. However, we also observe a birth year-linked increase in non-PGS variance in BMI (i.e., λ_1_ > 0), raising the possibility that the obtained GxE may be attributable to a more general pattern of increasing variance in BMI with birth year. In sex-pooled analyses, probabilities associated with tests of *H*_0_ : *ξ* = 0 are approximately 0.02. However, findings vary slightly by gender; especially for males, we are unable to reject the null of the scaling model. To illustrate the differences in implications in our sex-stratified analyses, we also conduct a posterior predictive check analysis [19], see Figure 2; there is less change in the penetrance of BMI PGS across birth years for males as compared to females. Using Eqn 43, we can decompose variation in *π*_1_ estimates. For females and males, 41% (95% CI: 10-57%) and 25% (95% CI: 0-54%) of the GxE (*π*_1_) effect cannot be attributed to a simple scaling term. In an analysis of all respondents, this proportion is 37% (95% CI: 9-52%). Collectively these results indicate substantial evidence for heteroscedasticity as a function of birth year; the simple homoscedastic GxE model does not represent the data adequately. Further discussion of this analysis and findings related to genetic heteroscedasticity are discussed in the SI.

**Figure 2:**
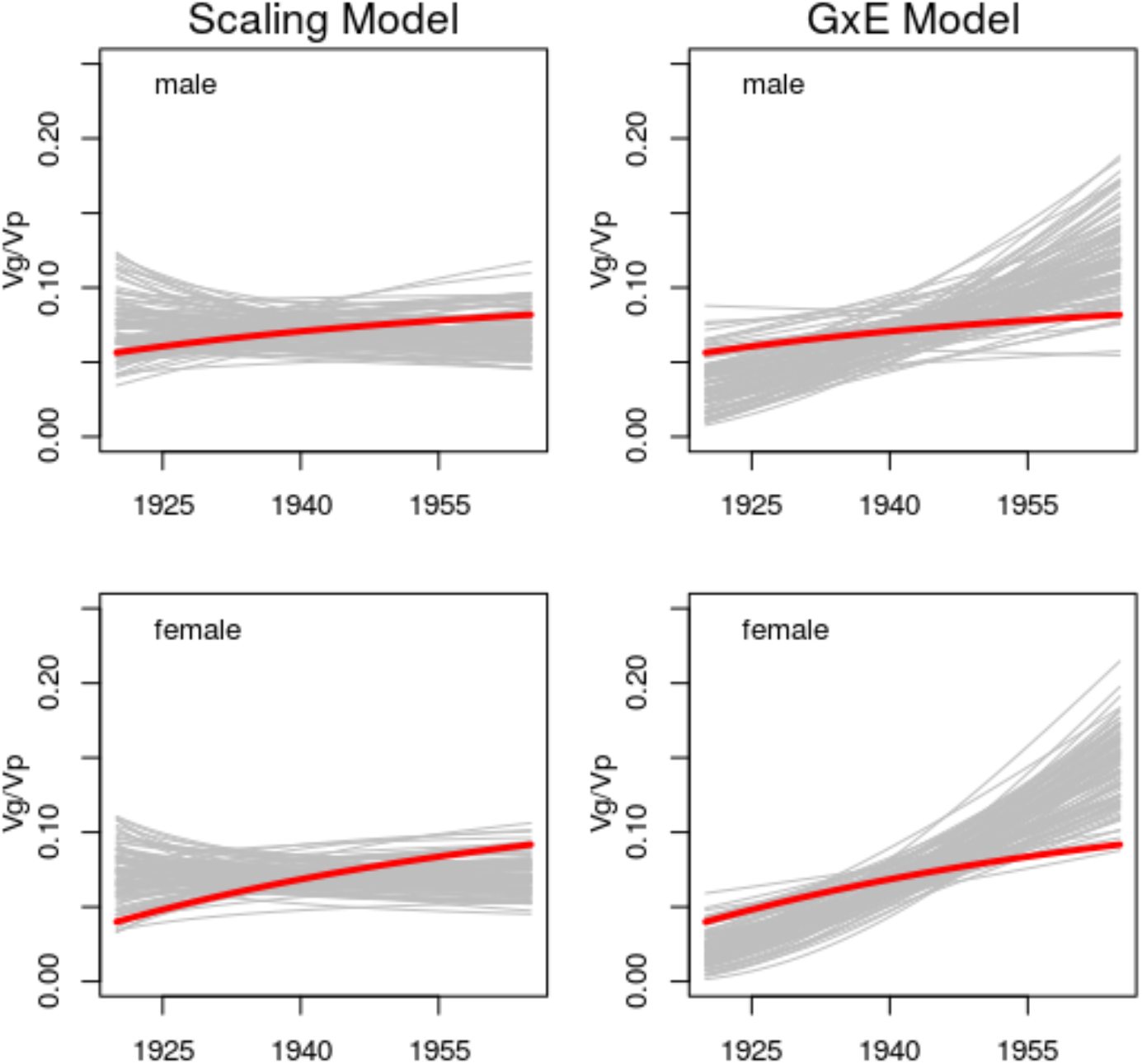
Gray lines represent genetic penetrance as a function of birth year simulated based on either the scaling model (Eqn 13, on left) or the standard homoscedastic GxE model (Eqn 1, on right). Red lines represent genetic penetrance as a function of birth year estimated from the real data using the heteroscedastic regression model (Eqn 8). Analyses based on standardized BMI data.

### 3.2 SNP Analyses

We conducted heteroscedastic regression analyses for 96 marker SNPs for the genome-wide significant loci identified in [16] using independent data from N=380,605 participants from UK Biobank (UKB; not used in the BMI GWAS [16]). Replication results for the main effects can be found in Table F.2. Note that the UKB is highly non-representative [20] potentially affecting the generalizability of our results.

The environmental heteroscedasticity parameter (λ_1_) was positive and significant in all models, suggesting greater variance in BMI among those born later in the 20th century. With the full heteroscedasticity model, we identified four SNPs that exhibited significant gene-by-birth year effects, as indicated by the *π*_1_ parameter at the Bonferroni-adjusted alpha level. Parameter estimates from the full heteroscedasticity model for these SNPs are reported in Table 1, along with *ξ_E_* and *ξ_G_* estimates obtained from the environmental and genetic heteroscedasticity models, respectively. Note that rs1558902 is a variant in the FTO gene locus. This finding replicates earlier results [21] which reported increasing penetrance of a variant with FTO over historical time using a different data set. However, *ξ_G_* was highly significant for each of these four SNPs, allowing us to reject the null that observed GxE is driven entirely by scaling.

**Table 1:** Estimates from parameters of the full heteroscedasticity model (Eqn 49) in analysis of GxE for Box-Cox transformed BMI as a function of birthyear in the UK Biobank for those SNPs with significant *π*_1_ estimates following Bonferonni adjustment. The *ξ_E_* and *ξ_G_* estimates reported are obtained from the environmental and genetic heteroscedasticity models, respectively. We show probabilities for parameters when the maximal probability in a column is larger than 1*e* – 6.

| SNP | $\pi_0$ | $\pi_1$ | $\text{Pr}_{\pi_1}$ | $\lambda_0$ | $\lambda_1$ | $\lambda_2$ | $\text{Pr}_{\lambda_2}$ | $\text{Pr}_{\xi_E}$ | $\text{Pr}_{\xi_G}$ |
| --- | --- | --- | --- | --- | --- | --- | --- | --- | --- |
| rs543874_A | -0.0271 | -0.0085 | 1.90e-07 | 0.9873 | 0.0369 | -0.0036 | 1.48e-03 | 4.39e-06 | 1.04e-07 |
| rs13021737_G | 0.0279 | 0.0062 | 1.33e-04 | 0.9873 | 0.0369 | 0.0036 | 1.53e-03 | 1.46e-03 | 1.75e-04 |
| rs1558902_T | -0.0503 | -0.0077 | 2.12e-06 | 0.9864 | 0.0368 | -0.0096 | 1.76e-17 | 2.96e-04 | 5.14e-07 |
| rs2075650_A | 0.0113 | -0.0060 | 2.43e-04 | 0.9875 | 0.0371 | -0.0002 | 8.53e-01 | 2.01e-04 | 5.78e-04 |

With the full heteroscedasticity model, we identified six SNPs that exhibited significant effects on the dispersion of BMI, as indicated by the λ_2_ parameter at the Bonferroni adjusted alpha level. Parameter estimates for these SNPs are reported in the supplement (Table F.1), along with *ξ_E_* and *ξ_G_* estimates obtained from the environmental and genetic heteroscedasticity models, respectively. Of the 4 Bonferonni significant variants for SNP-by-Year interaction described above, only the FTO variant (rs1558902) appears among the 6 Bonferonni significant variants for vQTL. This finding replicates earlier results [3] reporting that the FTO gene locus is associated with greater variability in BMI. The p value of *ξ_G_* for this variant is 5.1e-07, indicating the the above-reported SNP-by-birth-year interaction on BMI for FTO is not simply attributable to a general modulation of variance by rs1558902. Note that although the *ξ_G_* statistic is not significant for rs7903146, rs9925964, and rs2287019, these SNPs do not exhibit GxE at Bonferonni-corrected levels, rendering the *ξ_G_* moot.

To examine the total evidence of signal for GxE and genetic heteroscedasticity in the 96 SNPs, we provide a QQ plot of −1og10(p) for parameters indexing SNP-controlled heteroscedasticity of BMI vQTL and gene-by-birth year GxE interaction for Box-Cox transformed BMI in the left panel of Figure 3. We observe considerable departure of the observed −1og10(p) values relative to the expectation under the null, indicating strong enrichment of signal for both vQTL and GxE. In the right panel of Figure 3, we provide the scatter plots of parameter estimates for SNP main effects (*π*_0_), GxE (*π*_1_), for vQTL (λ_2_) for Box-Cox transformed BMI. We observe moderate-to-strong associations between the main effect of a SNP and both its GxE and vQTL effects, indicating that, among these variants selected on the basis of marking genome-wide significant loci for BMI in an external meta-analysis, those with larger main effects on BMI exhibit larger interactions with birth year and more pronounced moderation of BMI variance. Results of [6] suggest that similar patterns may apply genome-wide.

**Figure 3:**
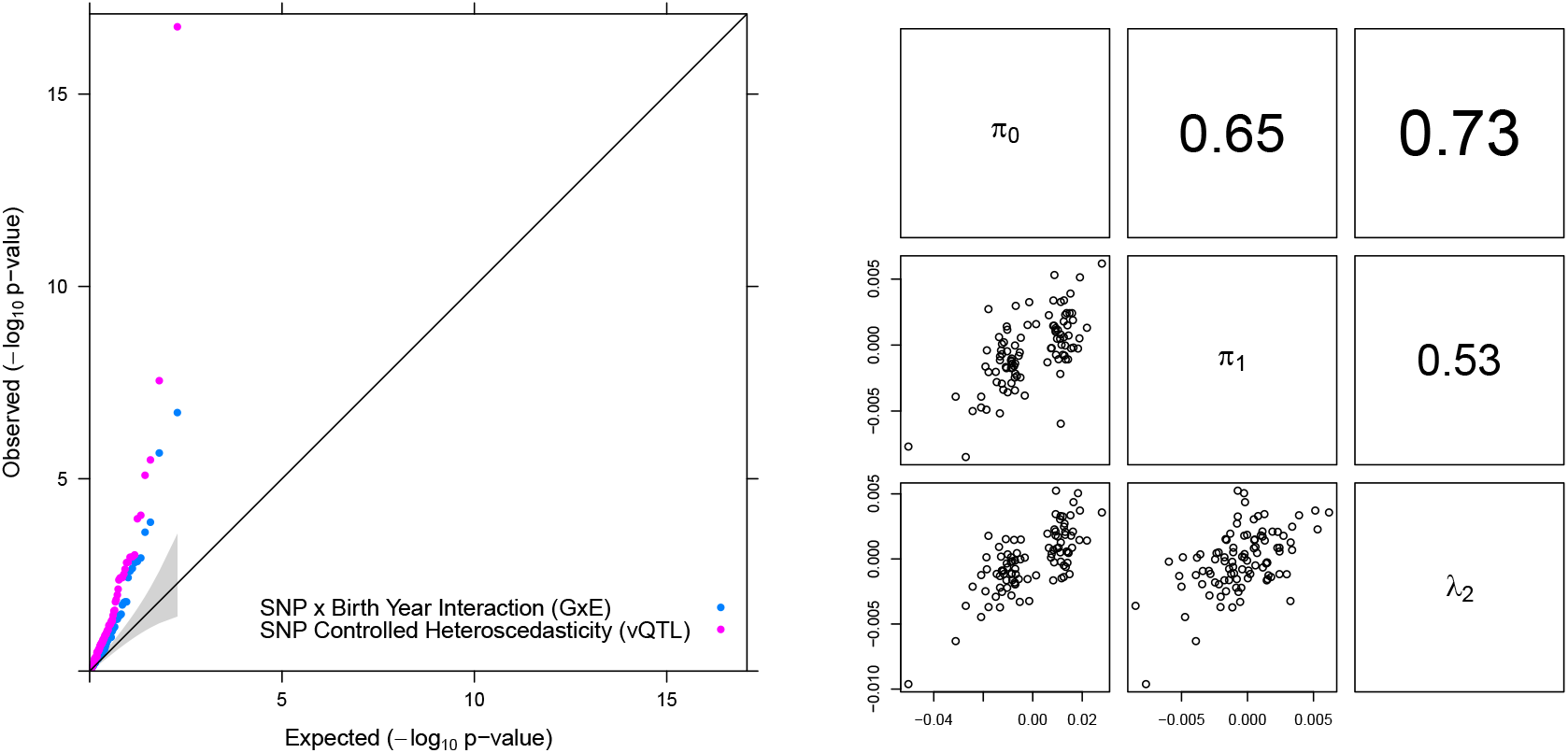
QQ plot of-1og10(p) for parameters indexing SNP-by-birth year interaction (GxE, as indicated by the *π*_1_ parameter) and SNP-controlled heteroscedasticity of BMI (vQTL, as indicated by the λ_2_ parameter) for Box-Cox transformed BMI. The considerable departure of the observed −1og10(p) values relative to the expectation under the null indicates strong enrichment of signal for both vQTL and GxE in these 96 SNPs previously associated with mean BMI in independent data [16]. Right: Multipanel Scatterplot of the parameters for the SNP main effect (*π*_0_), SNP-by-Birth Year interaction (*π*_1_), and vQTL (λ_2_) for Box-Cox transformed BMI. We analyzed 96 marker SNPs for the genome-wide significant loci for BMI identified in [16] in independent data.

Our results indicate that SNP moderation of birth year effects on BMI and SNP moderation of BMI dispersion were in the same direction as the SNP effects on BMI means. We observe considerable sign congruence between SNP main effects (*π*_0_) and SNP-by-Birth-Year interaction (*π*_0_) for 3 out of 4 SNPs exhibiting Bonferonni significant *π*_1_ estimates and for 71 out of 96 of all SNPs tested. We also observe considerable sign congruence between SNP main effects (*π*_0_) and vQTL effects (λ_2_) for 6 out of 6 SNPs exhibiting Bonferonni significant λ_2_ estimates, and for 75 out of 96 of all SNPs tested.

Finally, we note that the main effect of birth year (*τ*_1_) was approximately −0.058 in all models (all associated probabilities less than 1e-250), indicating that later-born UK Biobank participants tend to have lower BMI. Given substantial epidemiological evidence from representative samples indicating increasing BMI with birth year, we speculate that this negative association may be driven by selection bias in the UKB sample [20]. However, note that, despite decreasing mean BMI with birth year in this sample, we observe increasing variance in BMI with birth year (as indicated by the positive λ_1_ parameter), and increasing penetrance of SNPs with birth year (as indicated by strong sign congruence between *π*_0_ and *π*_1_ parameters). This further suggests that variance moderation and SNP-by-Birth Year interactions are unlikely to have resulted from positive associations between BMI mean and variance.

## 4 Discussion

When GxE is detected using the conventional regression-based model, it is common to interpret the interaction in terms of the specific genetic and environmental measures included in the interaction model. However, GxE may arise more generally when the genetic or environmental predictor moderates the total amount of variation in the phenotype. When the total amount of variation in a phenotype varies across the range of an environmental measure, traditional homoscedastic GxE models will detect interactions between that measure and all other correlates of the phenotype, both genetic and non-genetic. Here, we have delineated an expanded *heteroscedastic* GxE regression model that explicitly allows for heteroscedasticity in the phenotype as a function of the genetic and environmental measures. We use this model to derive a test statistic, *ξ*, which compares this heteroscedasticity model to a simpler *scaling model* in which GxE arises from more general differences in phenotypic variance, irrespective of etiology. When the scaling model holds, differences in phenotypic variance across the range of the genetic or environmental moderator induce a form of GxE that is only apparent in unstandardized units; the proportional contribution of the remaining predictors to phenotypic variance remains constant across the range of the moderator.

We provide an application of the heteroscedastic regression model to empirical data, replicating previous observations that that genetic predictors of BMI have become increasingly penetrant in recent years. However, we additionally observed that the total variance in BMI, including variation not associated with the genetic predictors, increased with participant birthyear. We found that, amongst males within the sample, the interaction between PGS and birth year was indistinguishable from a scaling model in which the total variance in BMI increases over birth year. For males, although the unstandardized variance in BMI accounted for by the PGS increased with birth year, the variance in BMI unaccounted for by the PGS also increased, such that the proportion of variance accounted for in BMI did not significantly vary with birth year. While we reject the null that the scaling model produced data for females, we still estimate that a large proportion of the observed GxE is due to scaling. Other work has considered the potential moderation of genetic risk for BMI as a function of diet [22, 23], other lifestyle characteristics [24], and educational attainment [25]. The issues discussed here may be a relevant feature of such empirical work; i.e., environmentally-linked heteroscedasticity may play a role in driving some previously observed GxE findings.

In instances in which the units and scaling of the phenotype are intrinsically-meaningful, biologically-meaningful, or policy-relevant, GxE that is observed on the unstandardized scale may be of high substantive and practical relevance. However, even when GxE arising from heteroscedasticity is deemed meaningful, it may still be important to discern whether interactions are specific to the variables selected for GxE or representative of a more general pattern that would be obtained for many correlates of the phenotype. It will also stil be important to understand whether the moderator is associated with differences in the percentile location of expected values along the distribution of the phenotype, or simply with differences in raw values (e.g. has a policy specifically increased the income of a particular subgroup, or increased the income of all subgroups, such that relative income of the subgroup of interest remains constant?). The heterosedastic regression models introduced here can be used to answer these questions.

Issues surrounding the scaling of the phenotype not fully explored here have the potential to have substantive effects on GxE results. For instance, skew in the distribution of the phenotype may produce both GxE and heteroscedasticity by inducing a dependency between the mean and the variance of the distribution of the phenotype. Such forms of GxE and may altogether dissappear under the appropriate tranformation, an issue that geneticists have be aware of since the days of Fischer [26]. Importantly, heteroscedasticity is not always simply an epiphenomenon of non-normality. For instance, in our empirical analysis of BMI, we report results of GxE analyses after a normalizing (Box-Cox) transform of BMI, and continue to detect both GxE and heteroscedasticity.

GxE research faces a number of conceptual and statistical challenges [8]. Thoughtfully and rigorously engaging with these challenges is particularly salient given the substantial recent increases in the availability of both genetic resources [27] and computational tools [28] for such research. Our proposed test is designed to help clarify the nature of potential GxE findings and to shed additional light on the processes contributing to variation in the phenotype; in particular, we can assess the extent to which the genetic etiology is relatively stable across the range of the environmental measure. Using the model and test statistic presented here, researchers can test for whether variance in the phenotype that is not explained by the genetic predictor shifts across the range of the environment and whether a scaling model may account for the obtained pattern of GxE. When the scaling model holds, we would suggest that the environment largely acts as a “dimming” mechanism on phenotypic variation [8] without altering the proportional contribution of genetic variation to the phenotype.

## Acknowledements

The authors would like to thank Dan Benjamin, Dalton Conley, Michel Nivard, Paul Rathouz, Mijke Rhemtulla, and Patrick Turley for helpful comments on an early version of this manuscript. This research was conducted using the UK Biobank Resource (Application No. 36046). This work was supported in part by the National Science Foundation Graduate Research Fellowship Program under Grant No. DGE-1656518 (ST), by the Institute of Education Sciences under Grant No. R305B140009 (ST), by NIH grants R01MH120219, R01AG054628, and R01HD083613 (EMTD), and by the Jacobs Foundation (EMTD). Any opinions expressed are those of the authors alone and should not be construed as representing the opinions of funding agencies. The authors declare no competing interests.

## Supplemental Information (SI)

### A Methods

#### A.1 Description of the Heteroscedastic Regression Model and Derivation of *ξ*

The *environmental heteroscedasticity model* is specified as

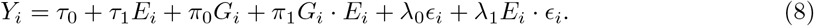

Here, *τ*_1_ is the main effect of the measured environment, *π*_0_ is the main effect of *G_i_*, *π*_0_ is the GxE directly analogous to *β*_3_ in Eqn 1, λ_0_ is the main effect of the error *ϵ_i_* (which we assume, without loss of generality given λ_0_, to have unit variance), and λ_1_ is the coefficient used to index heteroscedasticity.

This model will produce very similar unstandardized estimates of GxE as the standard GxE model given by Eqn 1. In other words, *π*_1_ in Eqn 8 is directly comparable to *β*_3_ in Eqn 1. However, the heteroscedasticity model further models whether variance in the phenotype that is not accounted for by the genetic predictor (*G_i_*) also varies as a function of the environment (*E_i_*) in the form of an *E_i_ϵ_i_* interaction, as captured by the λ_1_ parameter. Importantly, when *G_i_* only index portions of total genetic etiology of a trait [1–3], the *E_i_ϵ_i_* interaction may result from GxE with respect to the genetic factors not indexed by *G_i_*.

In order to test whether a more general pattern of heteroscedasticity as a function of *E_i_* is sufficient to account for the observed GxE, we propose a restricted form version of this model that does not directly model GxE but instead allows for the total variance of the phenotype to vary as a function of *E_i_*. This *scaling model* represents a scenario wherein GxE, in the sense of a significant estimate of *β*_3_ in Eqn 1, arises as a function of differences in the total variance of *Y* as a function of *E_i_*. The scaling model for moderation of the variance of *Y_i_* as a function of *E_i_* takes the form

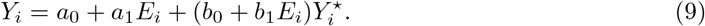

Here, *E_i_* is a measured environment and 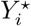 is an unobserved factor representing unexplained variation in *Y_i_* incremental of *E_i_*. We introduce a scaling transformation that produces heteroscedasticity in *Y_i_* as a function of *E_i_* via the *b*_0_ + *b*_1_*E*_i_ coefficient on the *Y*^⋆^ term. We let

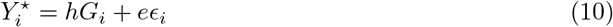

where *G* is a measured genotype standardized to have unit variance and *E_i_* is an unobserved error term (we assume it is also scaled to have unit variance). Finally, we set the scale of *Y*^⋆^ by specifying 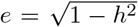. The penetrance of the measured gentoype is thus controlled via the relative magnitudes of *h* and *e*. Note, in particular, that in terms of 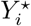, the penetrance of the measured genotype is constant. The test that we introduce relies on this property; i.e. that when properly scaled, the penetrance of the measured genotype to *Y_i_* is constant in this model. Suppose that *b*_1_ = 0. In that case, *Y_i_* is affected solely by environmental and genetic main effects. If *b*_1_ ≠ 0, then the role of 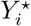 with respect to *Y_i_* varies as a function of *E*, and as we elucidate next, so do the raw (but not relative) contributions of *G_i_* and *ϵ_i_* to *Y*.

Substituting Eqn 10 into Eqn 9, and setting 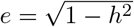, yields

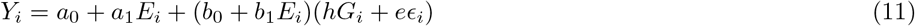

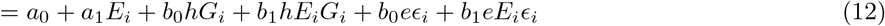

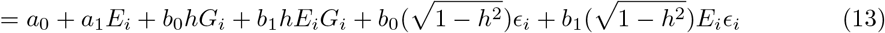

It can be seen that the environmental scaling model (Eqn 13) constitutes a constrained form of the environmental heteroscedasticity model (Eqn 8). Both equations take the same form in terms of *E* and *G*. However, whereas the coefficients in Eqn 8 are unconstrained, these coefficients are linked to one another in Eqn 13 via the the ratio *h/e* which, as noted above, is fixed under the scaling model. Thus, Eqn 8 is equivalent to Eqn 13 when

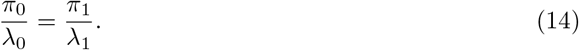

because, under Eqn 13

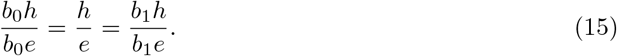

i.e.

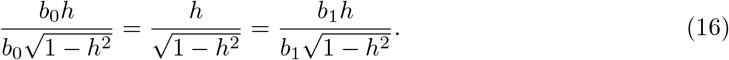

We use Eqn 14 as the basis for a hypothesis test. In particular, we consider the test statistic

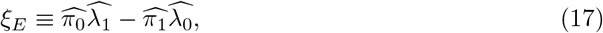

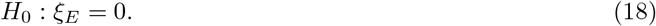

When *ξ_E_* = 0, the environmental heteroscedasticity model is indistinguishable from the simpler; scaling model, whereas when *ξ_E_* is differs from 0, the environmental scaling model is rejected.

#### A.2 Estimation

We estimate Eqn 8 using maximum likelihood by assuming *ϵ_i_* ~ Normal(0, 1).We assume that *Y*; is mean-centered and that both *G_i_* and *E_i_* are non-random. Consider 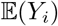 and 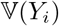. We first have

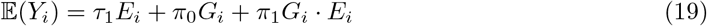

since 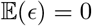. We then have

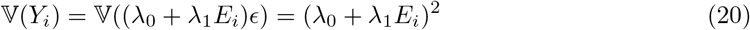

since we assume 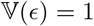. If we write

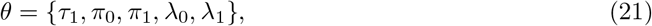

note that both 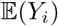 and 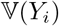 are functions of *θ*. Estimation of Eqn 8 is based on the likelihood,

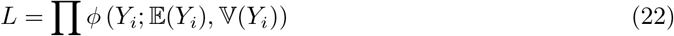

where *ϕ* is the normal density and we use the above expressions for 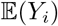 and 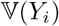. We can thus solve for θ = {τ_1_, *π*_0_, *π*_1_, λ_0_, λ_1_} using ML based on

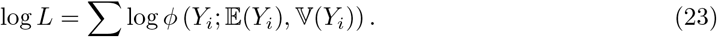

In particular, we obtain estimators via

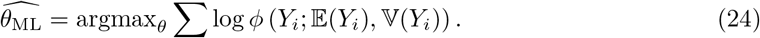

To obtain ML estimates, it is convenient to also have the gradient. Here, we have

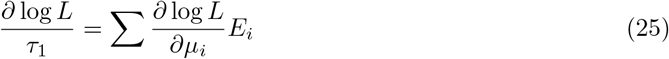

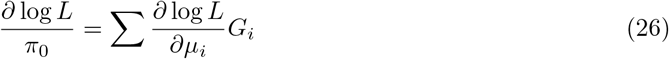

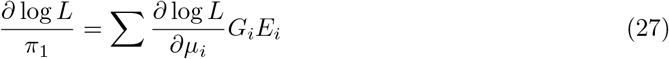

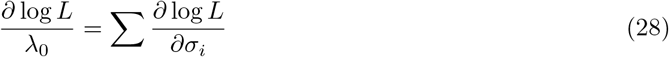

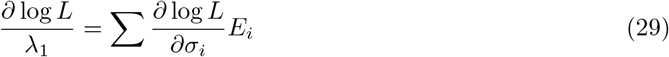

where 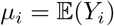, 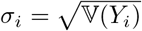, and

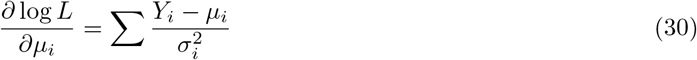

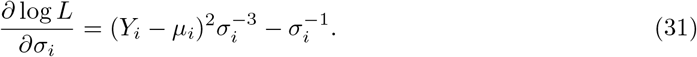

Using these estimates, we can compute a test statistic. Replacing the values in Eqn 15 with estimates from Eqn 8, we propose to test

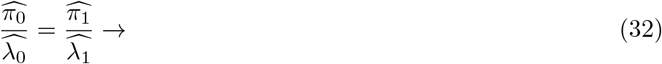

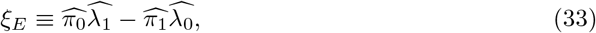

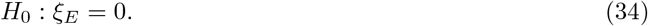

If *H*_0_ holds, the underlying form of GxE cannot be distinguished from the scaling phenomenon in which the total variance in *Y_i_* varies as a function of E. Alternatively put, failure to reject *H*_0_ constitutes a failure to reject the hypothesis that the *proportional* contributions of *G_i_* and *ϵ_i_* to *Y_i_* do not vary with E.

*H*_0_ can be tested via a Wald test for nonlinear restrictions. To test this nonlinear restriction, we consider (see Ref [4], Eqn 6–29)

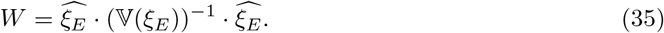

Given that we are testing a single restriction, W has a χ^2^ distribution with one degree of freedom in large samples. To estimate 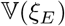, we use the Hessian of Eqn 23 (see B) and the delta method. Computationally, we do this via [5]. Note also that conventional tests of *π*_1_ and λ_1_ can be used to offer insight into the existence of GxE and potential heteroscedasticity; we make use of such tests below.

#### A.3 Extensions

##### A.3.1 Decomposition of *π*_1_

We can decompose the *π*_1_ estimate in Eqn 8 in a way that helps elucidate the contribution of heteroscedasticity to observed GxE. Consider an expanded version of the environmental scaling model (Eqn 9) wherein we add an explicit GxE term:

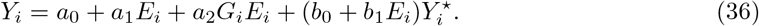

This expanded scaling model (in conjunction with Eqn 10) is equivalent (same number of free parameters and same model fit) to the environmental heteroscedasticy model, but decomposes the GxE effect into a component that is accounted for by scaling and a component unaccounted for by scaling (note that all environmental heteroscedasticity is assumed to operate through scaling). We can therefore equate Eqn 36 and Eqn 8 and recover the following system of equations:

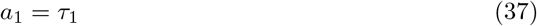

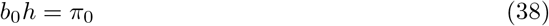

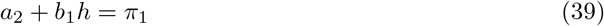

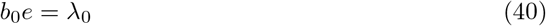

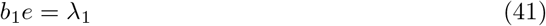

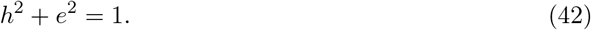

Using estimates from Eqn 8 for the quantitities on the right, we can solve for the parameters on the left. In particular, we can decompose *π*_1_ into *a*_2_ and *b*_1_*h*. The term *a*2 represents the degree to which observed data depart from the scaling model. Moreover, we can calculate the ratio

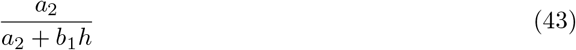

to describe the the magnitude of this departure as a proportion of the total GxE effect.

##### A.3.2 Genetically-linked heteroscedasticity

A version of the heteroscedasticity model presented in Eqn 8 above can be constructed so as to relax the assumption of homoscedasticity of residuals across *G_i_* as follows

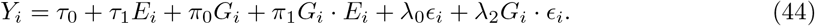

Here λ_2_ is the coefficient used to index heteroscedasticity as a function of *G_i_*. We assume homoscedasticity of residuals as a function of *E_i_* by omitting λ_1_. We refer to Eqn 44 as a *genetic heteroscedasticity model*. When a single variant is entered for *G*, this model consitutes a vQTL model that includes a term for GxE. Note that this model is equivalent to the model presented in Eqn 8, but with *G_i_* entered for *E_i_* and *E_i_* entered for *G_i_*.

The scaling model for moderation of the variance of *Y_i_* as a function of *G_i_* takes the form

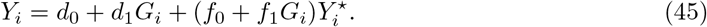

and

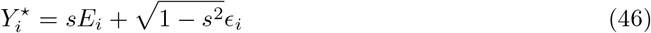

As in the scaling model for moderation the variance of *Y_i_* as a function of *E_i_*, *ϵ_i_* is assumed to have unit variance. In terms of 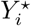, the effect of *E_i_* constant across *G_i_*. However, in terms of *Y_i_*, the effect of *E_i_* changes across *G_i_* as result of heteroscedasticity across *G_i_*.

It follows that the hypothesis test comparing the GxE model allowing for heterocedasticity as a function of *G_i_* (given by Eqn 44) to its reduced scaling model (given by Eqns 45 and 46, is

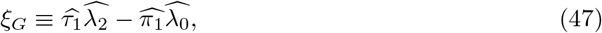

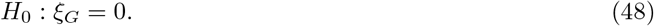

Finally, we can fit a *full heteroscedasticity model* that allows for moderation of residual variance in *Y_i_* as a function of both *G_i_* and *E_i_* as follows

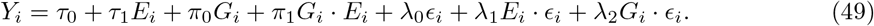

This model may be particularly useful when there is nonzero gene-environment correlation between the two moderators, i.e. | *r*(*G_i_,E_i_*) |> 0, such that *G_i_* and *E_i_* can be mutually controlled when modelling heteroscedasticity. However, to avoid added complexity, we recommend that comparisons with the scaling models rely on the *ξ_E_* and *ξ_G_* tests from the separate environmental heteroscedasticity and genetic heteroscedasticity models, respectively.

### B The Hessian

We use the Hessian 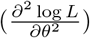 for construction of the covariance matrix of the estimation errors for the model parameters under the null model. The constituent parts of the Hessian are as follows.

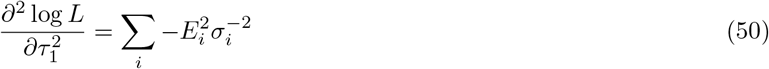

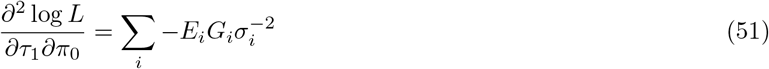

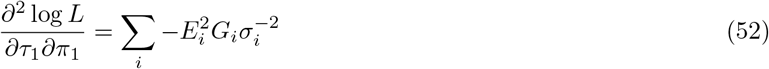

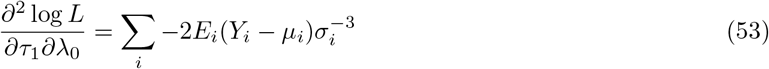

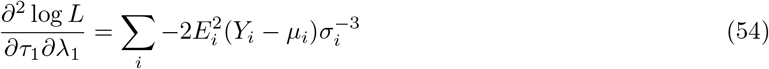

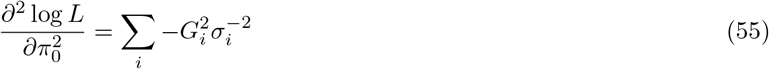

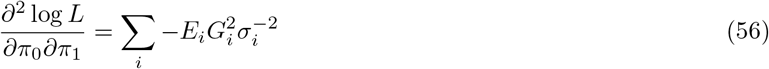

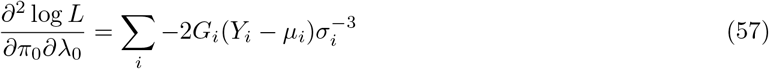

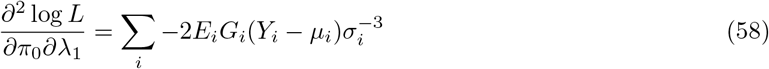

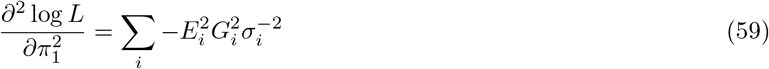

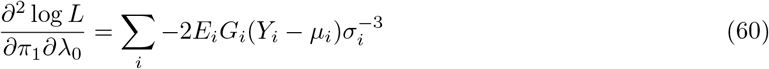

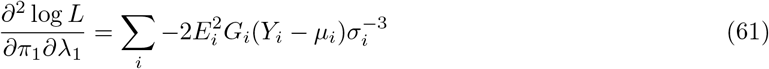

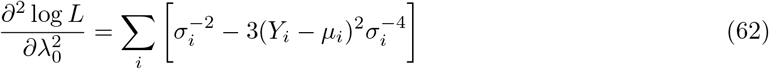

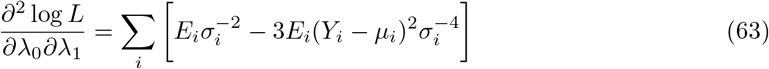

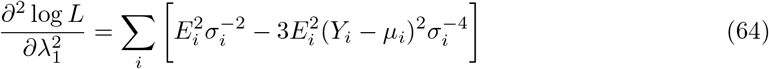

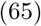

### C Simulation Study

We now consider both the null distribution of the test statistic under different configurations of the relevant parameters from the scaling model (Eqn 13) and the performance of our test statistic in the case of generating models that do not conform to the scaling model (i.e. when Eqn 1 is used to generate a model with GxE and homoscedastic residuals, or Eqn 8 is used to general a model with heteroscedastic residuals but no GxE). We primarily describe the simulation models in the context of the heteroscadasticity and scaling models in which the measured environment, *E_i_* moderates the residual variance of the phenotype, *Y_i_*, as given by Eqns 8 and 13. However, as described in A.3, these models are equivalent to those in which the measured phenotype, *G_i_* moderates the residual variance of *Y_i_* (after *G_i_* and *E_i_* are substituted for one another). Thus simulation results apply to both scenarios.

#### C.1 Distribution of the test statistic under *H*_0_

We first consider the behavior of the test statistic when the null hypothesis is true. We generate data according to the *scaling model* (Eqn 13), estimate the *heteroscedasticity model* (Eqn 8), and then calculate the *ξ_E_* test statistic and associated Wald-based probability. We start by illustrating that the appropriate transformation of *ξ_E_* has a χ^2^ distribution. To do this, we generate 10,000 datasets from the scaling model (Eqn 13) with *b*_0_ = 1, *b*_1_ = 0.15, *h* = 0.5, *N* = 5000. We compute *ξ_E_* and then transform it to *W* via Eqn 35. Results are in Figure C.1. The null distribution of *W* neatly aligns with the density of χ^2^ (1). Below, we report on *ξ_E_* given that it is a relatively intuitive function of the estimated parameters but also exploit the fact that *ξ_E_* can be mapped to W and inference can be done via reference to χ^2^(1).

**Figure C.1:**
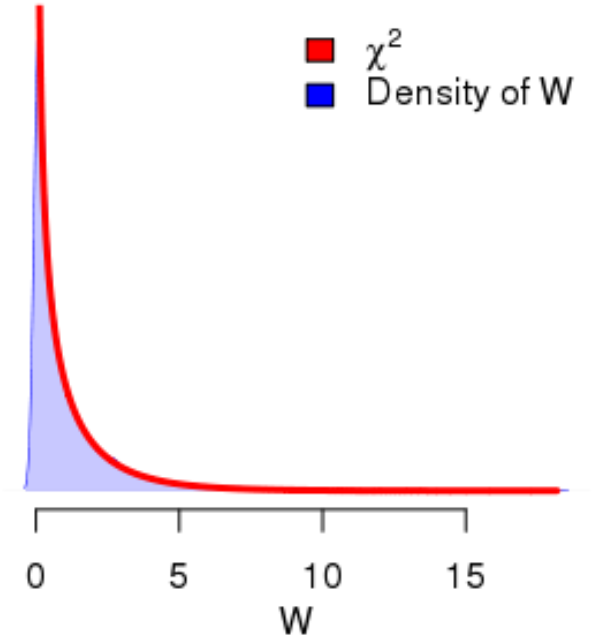
Comparison of distribution of W under the null hypothesis to χ^2^(1).

We next considered the behavior of *ξ_E_* under the null, for configurations of the relevant parameters in the scaling model (Eqn 13) meant to capture the relevant features of an analysis in which *G* is a PGS. We considered variation in four parameters. We used three different sample sizes (1000, 5000, 10000). We used two values respectively for *b*_0_ (0.8, 1) and *b*_1_ (0.05, 0.1). Finally, we used two values for *h* (0.1 and 0.5). The values of h suggest weakly (0.1) to strongly (0.5) penetrant genetic predictors; they were chosen to represent the range of potential PGS predictors given existing technology [1]. Recall that the value of *E* is fixed as a function of h. Note that we considered two different distributions for *E*; we sampled *E* from both the standard normal and the uniform distributions (where the uniform distribution was scaled to have the same standard deviation as the standard normal). As polygenic scores for complex traits are expected to be normally distributed [6], *G* was sampled from the standard normal distribution. To help develop intuition, suppose *b*_0_ = 0.8, *b*_1_ = 0.05, and *h* = 0.5. In that case, 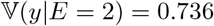 compared to 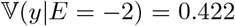, a ratio of 1.74 (i.e., there is heteroscedasticity). Finally, it is important to note that the scale of *ξ_E_* is dependent on several features (e.g. scale of *G* and *E*); patterns shown here are informative about the behavior of *ξ_E_* but the values themselves do not necessarily generalize to other cases (e.g., if *E* or *G* does not have unit variance). In contrast, W and associated tests are invariant to scaling of *G* and *E*, but sensitive to sample size.

Results are shown in Table C.1. Note that the estimates of *ξ_E_* are unbiased (i.e., the means are nearly zero). Further, as we would anticipate, the false positive rate (FPR) of the test is insensitive to the parameters; note also that the mean probability associated with the test of *ξ_E_* via Eqn 35 is near 0.05, as we would expect when the null is true.^1^

In contrast, the variance of the test statistic (and, thus, statistical power) depends on the sample size and simulation parameters. In particular, note the decline in the SD of the test statistics as a function of the sample size. Statistical power will also depend upon h; this test will have less power with less penetrant genetic predictors.

**Table C.1:**
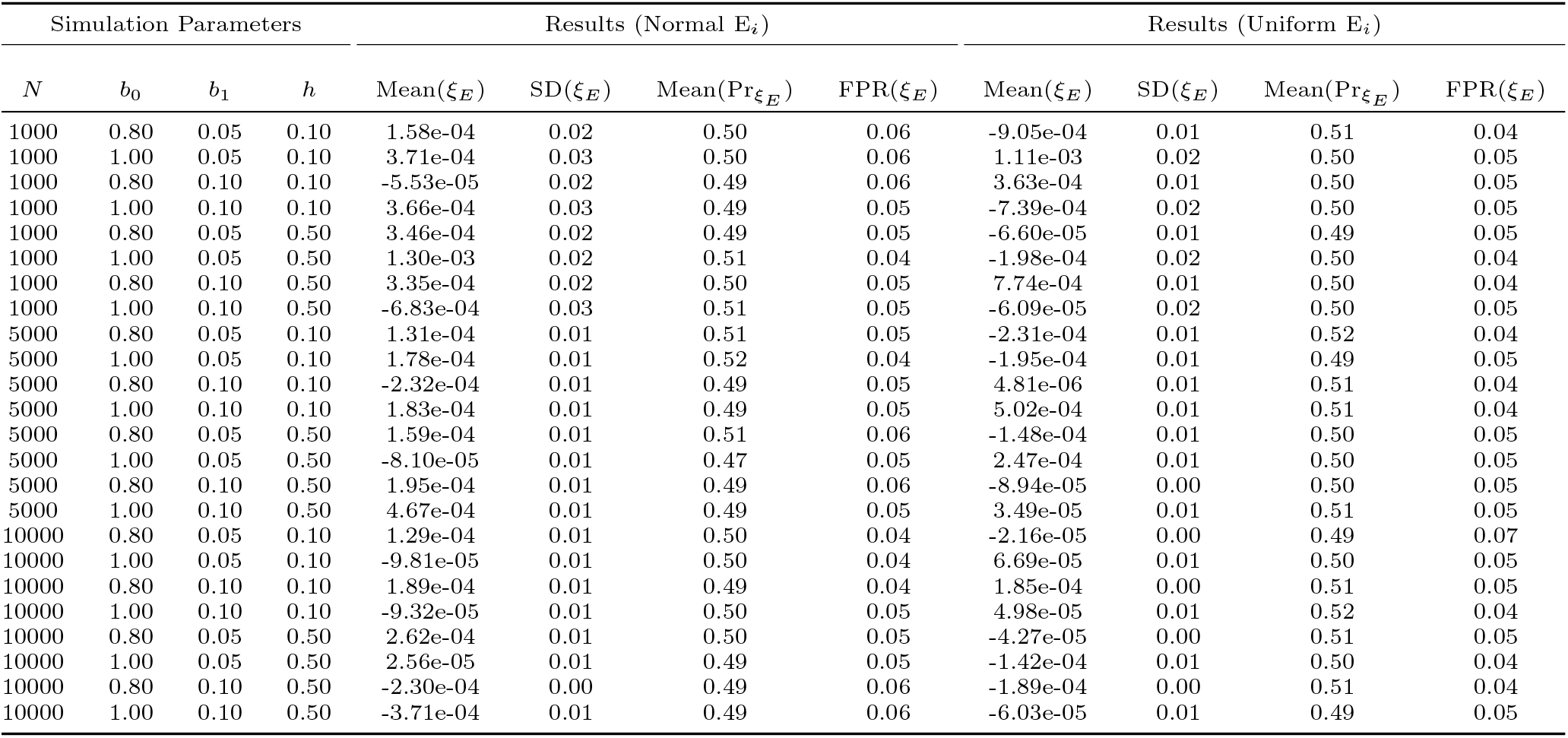
Distribution of *ξ_E_* under the null (mean and SD) and associated test (Eqn 35) for different values of parameters in Eqn 13. For each set of parameters, 1000 datasets are simulated. Normal and Uniform refer to the distributions used to generate the environmental variable.

We next considered the behavior of of *ξ_E_* under the null for configurations of parameters meant to capture the relevant features of an analysis in which *G* is an allele count for a single locus (e.g. a SNP). Results are in Table C.2. The table is similar to Table C.1, but where we now consider allele counts based on variants simulated under a binomial distribution with different minor allele frequencies (denoted p). Based on Extended Data Figure 2 in [7], we chose effect sizes of *h*^2^ ∈ {.05%, .1%}. Note that after generating *G* under a binomial distribution, it was standardized, such that h can be interpreted as a standardized effect size. We examine behavior using a sample size of *N* = 10000, which might be used for confirmatory tests of a select number of variants previously identified in a larger independent GWAS, and a sample size of *N* = 1000000, which might be used for genome-wide discovery. Simulation results indicate that the heteroscedasticity model and the *ξ_E_* statistic behave as expected across the range of conditions examined.

#### C.2 Performance under alternatives

We now turn to study the behavior of the heteroscedasticity model and associated *ξ_E_* test statistic when the null hypothesis is false (i.e., when Eqn 13 is not the data generating model). We consider two different data generating models: (i) a model with GxE and homoscedastic residuals, generated according to Eqn 1 (ii) a model with no GxE but with heteroscedastic residuals, generated according to Eqn 8. In these analyses, we aim to probe whether the test statistic allows us to reject Ho under different configurations of the relevant data-generating parameters.

**Table C.2:** Distribution of *ξ_E_* under the null (mean and SD) and associated test (via Eqn 35) for different values of parameters in Eqn 13 based on a standardized single-locus allele count. For each set of parameters, 100 datasets are simulated. The environmental variable is sampled from the standard normal distribution.

| $N$ | $b_0$ | $b_1$ | $h$ | $MAF$ | $\widehat{\pi}_1$ | $\widehat{\lambda}_1$ | Mean($\xi_E$) | SD($\xi_E$) | Mean($\text{Pr}_{\xi_E}$) | $\frac{\mathbb{E}(SE(\xi_E))}{SD(\xi_E)}$ |
| --- | --- | --- | --- | --- | --- | --- | --- | --- | --- | --- |
| 10000 | 0.80 | 0.05 | 0.02 | 0.01 | 6.91e-04 | 0.05 | 3.25e-04 | 6.50e-03 | 0.51 | 0.97 |
| 10000 | 1.00 | 0.05 | 0.02 | 0.01 | 6.52e-04 | 0.05 | 4.67e-04 | 9.76e-03 | 0.49 | 1.02 |
| 10000 | 0.80 | 0.10 | 0.02 | 0.01 | 3.25e-03 | 0.10 | -7.18e-04 | 6.35e-03 | 0.47 | 0.95 |
| 10000 | 1.00 | 0.10 | 0.02 | 0.01 | 2.53e-03 | 0.10 | -3.72e-04 | 9.29e-03 | 0.52 | 1.05 |
| 10000 | 0.80 | 0.05 | 0.03 | 0.01 | 1.17e-03 | 0.05 | 3.92e-04 | 6.18e-03 | 0.52 | 1.02 |
| 10000 | 1.00 | 0.05 | 0.03 | 0.01 | 1.81e-03 | 0.05 | -1.64e-04 | 1.13e-02 | 0.47 | 0.88 |
| 10000 | 0.80 | 0.10 | 0.03 | 0.01 | 2.29e-03 | 0.10 | 8.04e-04 | 6.13e-03 | 0.48 | 0.98 |
| 10000 | 1.00 | 0.10 | 0.03 | 0.01 | 2.43e-03 | 0.10 | 7.48e-04 | 9.56e-03 | 0.49 | 1.01 |
| 100000 | 0.80 | 0.05 | 0.02 | 0.01 | 1.27e-03 | 0.05 | -1.12e-04 | 1.96e-03 | 0.52 | 1.02 |
| 100000 | 1.00 | 0.05 | 0.02 | 0.01 | 6.32e-04 | 0.05 | 4.75e-04 | 3.10e-03 | 0.47 | 1.01 |
| 100000 | 0.80 | 0.10 | 0.02 | 0.01 | 2.00e-03 | 0.10 | 1.59e-04 | 1.84e-03 | 0.52 | 1.03 |
| 100000 | 1.00 | 0.10 | 0.02 | 0.01 | 2.49e-03 | 0.10 | -2.23e-04 | 2.97e-03 | 0.50 | 1.02 |
| 100000 | 0.80 | 0.05 | 0.03 | 0.01 | 1.21e-03 | 0.05 | 2.92e-04 | 2.11e-03 | 0.47 | 0.94 |
| 100000 | 1.00 | 0.05 | 0.03 | 0.01 | 1.40e-03 | 0.05 | 1.70e-04 | 2.88e-03 | 0.52 | 1.09 |
| 100000 | 0.80 | 0.10 | 0.03 | 0.01 | 3.04e-03 | 0.10 | 7.05e-05 | 2.16e-03 | 0.46 | 0.88 |
| 100000 | 1.00 | 0.10 | 0.03 | 0.01 | 2.53e-03 | 0.10 | 5.93e-04 | 3.04e-03 | 0.53 | 1.00 |
| 10000 | 0.80 | 0.05 | 0.02 | 0.05 | 3.28e-03 | 0.05 | -1.73e-03 | 5.97e-03 | 0.52 | 1.06 |
| 10000 | 1.00 | 0.05 | 0.02 | 0.05 | 1.03e-03 | 0.05 | 9.44e-05 | 1.15e-02 | 0.41 | 0.86 |
| 10000 | 0.80 | 0.10 | 0.02 | 0.05 | 2.20e-03 | 0.10 | 1.36e-04 | 5.53e-03 | 0.53 | 1.08 |
| 10000 | 1.00 | 0.10 | 0.02 | 0.05 | 2.61e-03 | 0.10 | -8.78e-06 | 1.04e-02 | 0.47 | 0.93 |
| 10000 | 0.80 | 0.05 | 0.03 | 0.05 | 7.91e-04 | 0.05 | 6.14e-04 | 6.44e-03 | 0.47 | 0.98 |
| 10000 | 1.00 | 0.05 | 0.03 | 0.05 | 1.51e-03 | 0.05 | -4.78e-06 | 1.13e-02 | 0.47 | 0.88 |
| 10000 | 0.80 | 0.10 | 0.03 | 0.05 | 4.31e-03 | 0.10 | -7.83e-04 | 6.14e-03 | 0.50 | 0.98 |
| 10000 | 1.00 | 0.10 | 0.03 | 0.05 | 4.16e-03 | 0.10 | -9.08e-04 | 9.64e-03 | 0.48 | 0.99 |
| 100000 | 0.80 | 0.05 | 0.02 | 0.05 | 1.16e-03 | 0.05 | -5.90e-05 | 1.98e-03 | 0.54 | 1.00 |
| 100000 | 1.00 | 0.05 | 0.02 | 0.05 | 7.56e-04 | 0.05 | 3.44e-04 | 3.06e-03 | 0.49 | 1.02 |
| 100000 | 0.80 | 0.10 | 0.02 | 0.05 | 1.74e-03 | 0.10 | 3.91e-04 | 2.03e-03 | 0.48 | 0.93 |
| 100000 | 1.00 | 0.10 | 0.02 | 0.05 | 2.20e-03 | 0.10 | 3.24e-05 | 3.20e-03 | 0.47 | 0.95 |
| 100000 | 0.80 | 0.05 | 0.03 | 0.05 | 1.72e-03 | 0.05 | -9.93e-05 | 1.83e-03 | 0.54 | 1.09 |
| 100000 | 1.00 | 0.05 | 0.03 | 0.05 | 1.31e-03 | 0.05 | 2.68e-04 | 3.31e-03 | 0.46 | 0.95 |
| 100000 | 0.80 | 0.10 | 0.03 | 0.05 | 3.69e-03 | 0.10 | -3.60e-04 | 2.15e-03 | 0.44 | 0.88 |
| 100000 | 1.00 | 0.10 | 0.03 | 0.05 | 3.09e-03 | 0.10 | 5.50e-05 | 3.35e-03 | 0.45 | 0.91 |
| 10000 | 0.80 | 0.05 | 0.02 | 0.50 | -4.27e-04 | 0.05 | 1.22e-03 | 6.81e-03 | 0.48 | 0.92 |
| 10000 | 1.00 | 0.05 | 0.02 | 0.50 | 1.53e-03 | 0.05 | -4.68e-04 | 8.87e-03 | 0.53 | 1.11 |
| 10000 | 0.80 | 0.10 | 0.02 | 0.50 | 1.60e-03 | 0.10 | 5.75e-04 | 6.84e-03 | 0.45 | 0.87 |
| 10000 | 1.00 | 0.10 | 0.02 | 0.50 | 3.90e-03 | 0.10 | -1.55e-03 | 9.61e-03 | 0.53 | 0.99 |
| 10000 | 0.80 | 0.05 | 0.03 | 0.50 | 1.35e-03 | 0.05 | 1.87e-04 | 6.44e-03 | 0.51 | 0.98 |
| 10000 | 1.00 | 0.05 | 0.03 | 0.50 | 3.36e-03 | 0.05 | -1.68e-03 | 9.08e-03 | 0.56 | 1.08 |
| 10000 | 0.80 | 0.10 | 0.03 | 0.50 | 4.00e-03 | 0.10 | -6.84e-04 | 5.63e-03 | 0.52 | 1.06 |
| 10000 | 1.00 | 0.10 | 0.03 | 0.50 | 3.78e-03 | 0.10 | -5.68e-04 | 9.75e-03 | 0.50 | 0.99 |
| 100000 | 0.80 | 0.05 | 0.02 | 0.50 | 1.34e-03 | 0.05 | -1.60e-04 | 2.00e-03 | 0.48 | 1.00 |
| 100000 | 1.00 | 0.05 | 0.02 | 0.50 | 7.07e-04 | 0.05 | 4.14e-04 | 2.91e-03 | 0.51 | 1.08 |
| 100000 | 0.80 | 0.10 | 0.02 | 0.50 | 2.21e-03 | 0.10 | 3.19e-05 | 1.92e-03 | 0.51 | 0.99 |
| 100000 | 1.00 | 0.10 | 0.02 | 0.50 | 2.00e-03 | 0.10 | 2.02e-04 | 2.73e-03 | 0.55 | 1.11 |
| 100000 | 0.80 | 0.05 | 0.03 | 0.50 | 1.89e-03 | 0.05 | -2.42e-04 | 2.24e-03 | 0.44 | 0.89 |
| 100000 | 1.00 | 0.05 | 0.03 | 0.50 | 1.99e-03 | 0.05 | -4.30e-04 | 3.26e-03 | 0.52 | 0.96 |
| 100000 | 0.80 | 0.10 | 0.03 | 0.50 | 3.00e-03 | 0.10 | 1.23e-04 | 2.04e-03 | 0.44 | 0.92 |
| 100000 | 1.00 | 0.10 | 0.03 | 0.50 | 3.51e-03 | 0.10 | -3.25e-04 | 2.96e-03 | 0.50 | 1.02 |

We first study a circumstance in which the generating model is GxE with homoscedastic residuals. We use Eqn 1 for our data-generating model and suppose that *e_i_* is white-noise (i.e., *e_i_* ~ N(0, *σ*^2^)) as opposed to heteroscedastic as a function of *E_i_*. In particular, we set *σ*^2^ = 1. We focus on two tests: a test of whether *π*_1_ =0 and a test of whether *ξ_E_* = 0. Data are analyzed with the environmental heteroscadasticity model (Eqn 8, which subsumes Eqn 1).

Results are shown in Table C.3. When both *N* and *b*_3_ are relatively small, we are poorly powered to detect GxE and to reject the scaling hypothesis (i.e., both Pr_π_1__ and Pr_*ξ_E_*_ are relatively large). Importantly, when GxE is not detected, the *ξ_E_* test is moot. In contrast, power to detect GxE and to reject scaling are both excellent when *N* or *b*_3_ is relatively large. Throughout we reject λ_1_ = 0 for *α* = .05 at the appropriate FPR.

**Table C.3:** Behavior of *ξ_E_* when true model is homoscedastic GxE (Eqn 1). Behavior is shown for 1000 simulated datasets based on each set of parameters.

| Simulation Parameters |  |  |  | Results |  |  |  |  |
| --- | --- | --- | --- | --- | --- | --- | --- | --- |
| $N$ | $\beta_1$ | $\beta_2$ | $\beta_3$ | Mean($\Pr_{\pi_1}$) | Power($\pi_1$) | Mean($\Pr_{\xi_E}$) | Power($\xi_E$) | FPR($\lambda_1$) |
| 1000 | 0.10 | 0.00 | 0.05 | 2.45e-01 | 3.48e-01 | 2.48e-01 | 3.48e-01 | 0.05 |
| 1000 | 0.20 | 0.00 | 0.05 | 2.26e-01 | 3.33e-01 | 2.31e-01 | 3.35e-01 | 0.05 |
| 1000 | 0.10 | 0.20 | 0.05 | 2.36e-01 | 3.56e-01 | 2.38e-01 | 3.57e-01 | 0.07 |
| 1000 | 0.20 | 0.20 | 0.05 | 2.16e-01 | 3.87e-01 | 2.21e-01 | 3.80e-01 | 0.05 |
| 1000 | 0.10 | 0.50 | 0.05 | 2.40e-01 | 3.34e-01 | 2.41e-01 | 3.32e-01 | 0.05 |
| 1000 | 0.20 | 0.50 | 0.05 | 2.23e-01 | 3.57e-01 | 2.26e-01 | 3.41e-01 | 0.06 |
| 1000 | 0.10 | 0.00 | 0.15 | 7.68e-04 | 9.98e-01 | 8.40e-04 | 9.97e-01 | 0.05 |
| 1000 | 0.20 | 0.00 | 0.15 | 9.08e-04 | 9.97e-01 | 1.01e-03 | 9.95e-01 | 0.04 |
| 1000 | 0.10 | 0.20 | 0.15 | 8.63e-04 | 9.98e-01 | 8.68e-04 | 9.97e-01 | 0.06 |
| 1000 | 0.20 | 0.20 | 0.15 | 6.13e-04 | 9.97e-01 | 6.72e-04 | 9.98e-01 | 0.05 |
| 1000 | 0.10 | 0.50 | 0.15 | 8.49e-04 | 9.96e-01 | 9.04e-04 | 9.97e-01 | 0.06 |
| 1000 | 0.20 | 0.50 | 0.15 | 9.02e-04 | 9.97e-01 | 1.14e-03 | 9.95e-01 | 0.06 |
| 5000 | 0.10 | 0.00 | 0.05 | 1.30e-02 | 9.37e-01 | 1.30e-02 | 9.32e-01 | 0.04 |
| 5000 | 0.20 | 0.00 | 0.05 | 9.69e-03 | 9.53e-01 | 1.03e-02 | 9.47e-01 | 0.06 |
| 5000 | 0.10 | 0.20 | 0.05 | 1.39e-02 | 9.37e-01 | 1.39e-02 | 9.32e-01 | 0.05 |
| 5000 | 0.20 | 0.20 | 0.05 | 1.26e-02 | 9.34e-01 | 1.34e-02 | 9.30e-01 | 0.05 |
| 5000 | 0.10 | 0.50 | 0.05 | 1.29e-02 | 9.43e-01 | 1.29e-02 | 9.43e-01 | 0.04 |
| 5000 | 0.20 | 0.50 | 0.05 | 1.52e-02 | 9.36e-01 | 1.60e-02 | 9.36e-01 | 0.05 |
| 5000 | 0.10 | 0.00 | 0.15 | 3.05e-13 | 1.00e+00 | 1.36e-12 | 1.00e+00 | 0.06 |
| 5000 | 0.20 | 0.00 | 0.15 | 1.62e-17 | 1.00e+00 | 9.32e-17 | 1.00e+00 | 0.04 |
| 5000 | 0.10 | 0.20 | 0.15 | 1.58e-15 | 1.00e+00 | 1.16e-15 | 1.00e+00 | 0.05 |
| 5000 | 0.20 | 0.20 | 0.15 | 2.74e-17 | 1.00e+00 | 5.83e-17 | 1.00e+00 | 0.04 |
| 5000 | 0.10 | 0.50 | 0.15 | 7.99e-18 | 1.00e+00 | 2.30e-17 | 1.00e+00 | 0.05 |
| 5000 | 0.20 | 0.50 | 0.15 | 1.48e-14 | 1.00e+00 | 2.87e-14 | 1.00e+00 | 0.05 |

We next consider the circumstance in which there is no GxE, but there are *G* and *E* main effects, and there is heteroscedasticity as a function of *E* so as to parallel Eqn 20. In particular, we generate data via

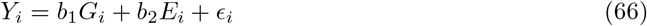

with *ϵ_i_* ~ N(0, (λ_0_ + λ_1_*E_i_*)^2^). We use λ_0_ = 2 and vary λ_1_ between 0.15 and 0.3 (if λ_1_ = 0, there is no heteroscedasticity). We fit the environmental heteroscedasticity model (eqn 8) and focus on tests of *π*_1_ =0 (which is true under the generating model), λ_1_ =0 (which is false under the generating model), and *ξ_E_* =0 (which is false under the generating model).

Results are shown in Table C.4. First, note that the appropriate FPR of .05 is obtained for *π*_1_ = 0 at alpha=.05, irrespective of λ_0_ and λ_1_. This indicates that heteroscedasticity has not induced the impression of GxE under these conditions. We can also uniformly detect that λ_1_ =0 suggesting that errors are in fact heteroscedastic. However, tests of *ξ_E_* are relatively weak here (e.g., power is 0.2 at best). Importantly, as the tests of *π*_1_ appropriate indicate no GxE, the question of whether there is scaling GxE is moot.

**Table C.4:** Behavior of *ξ_E_* when true model does not have GxE but does have heteroscedastic error (Eqn 66). Behavior is shown for 1000 simulated datasets based on each set of parameters.

| Simulation Parameters |  |  |  |  | Results |  |  |  |  |  |
| --- | --- | --- | --- | --- | --- | --- | --- | --- | --- | --- |
| $N$ | $\pi_0$ | $\tau_1$ | $\lambda_0$ | $\lambda_1$ | Mean($\text{Pr}_{\pi_1}$) | FPR($\pi_1$) | Mean($\text{Pr}_{\lambda_1}$) | Power($\lambda_1$) | Mean($\text{Pr}_{\xi_E}$) | Power($\xi_E$) |
| 1000.00 | 0.10 | 0.00 | 2.00 | 0.15 | 0.50 | 0.06 | 2.03e-02 | 0.90 | 0.49 | 0.05 |
| 1000.00 | 0.20 | 0.00 | 2.00 | 0.15 | 0.50 | 0.05 | 1.55e-02 | 0.93 | 0.49 | 0.06 |
| 1000.00 | 0.10 | 0.50 | 2.00 | 0.15 | 0.49 | 0.07 | 1.74e-02 | 0.92 | 0.48 | 0.07 |
| 1000.00 | 0.20 | 0.50 | 2.00 | 0.15 | 0.50 | 0.05 | 1.70e-02 | 0.93 | 0.48 | 0.06 |
| 1000.00 | 0.10 | 0.00 | 2.00 | 0.30 | 0.50 | 0.05 | 1.62e-06 | 1.00 | 0.48 | 0.05 |
| 1000.00 | 0.20 | 0.00 | 2.00 | 0.30 | 0.50 | 0.05 | 8.32e-07 | 1.00 | 0.45 | 0.09 |
| 1000.00 | 0.10 | 0.50 | 2.00 | 0.30 | 0.50 | 0.06 | 5.46e-06 | 1.00 | 0.49 | 0.07 |
| 1000.00 | 0.20 | 0.50 | 2.00 | 0.30 | 0.49 | 0.05 | 8.40e-07 | 1.00 | 0.46 | 0.07 |
| 5000.00 | 0.10 | 0.00 | 2.00 | 0.15 | 0.51 | 0.06 | 1.25e-08 | 1.00 | 0.51 | 0.06 |
| 5000.00 | 0.20 | 0.00 | 2.00 | 0.15 | 0.49 | 0.05 | 9.85e-08 | 1.00 | 0.46 | 0.09 |
| 5000.00 | 0.10 | 0.50 | 2.00 | 0.15 | 0.49 | 0.06 | 6.29e-09 | 1.00 | 0.48 | 0.06 |
| 5000.00 | 0.20 | 0.50 | 2.00 | 0.15 | 0.50 | 0.05 | 5.07e-08 | 1.00 | 0.46 | 0.08 |
| 5000.00 | 0.10 | 0.00 | 2.00 | 0.30 | 0.49 | 0.06 | 6.79e-36 | 1.00 | 0.45 | 0.10 |
| 5000.00 | 0.20 | 0.00 | 2.00 | 0.30 | 0.50 | 0.06 | 1.95e-37 | 1.00 | 0.33 | 0.22 |
| 5000.00 | 0.10 | 0.50 | 2.00 | 0.30 | 0.49 | 0.05 | 7.32e-38 | 1.00 | 0.44 | 0.09 |
| 5000.00 | 0.20 | 0.50 | 2.00 | 0.30 | 0.49 | 0.06 | 1.48e-38 | 1.00 | 0.32 | 0.23 |

### D Software

Code to estimate the heteroscestacity model and to estimate *ξ_E_* is available in an R package.^2^ In particular, the documentation illustrates functionality via simple simulations similar to those found in Section C.

### E Data

#### E.1 HRS Data

The first wave of HRS data collection was in 1992, the most recent in 2018 (wave 13). We focus on N=11,586 respondents of European ancestry. Descriptive information for the environment and outcome are shown in Figure E.1. We use data from the RAND HRS files on BMI, see [8] for additional documentation. In particular, we consider the mean BMI over all waves. The mean BMI was skewed, so we also considered a transformed version (see Figure E.1). HRS respondents were born across a wide span of birthyears centered around 1940. To ensure that results aren’t driven by small numbers of observations with extreme birth years, we analyzed those born between 1920–1965. All data (polygenic score, birthyear, and BMI) are standardized to have mean zero and unit variance for analysis.

**Figure E.1:**
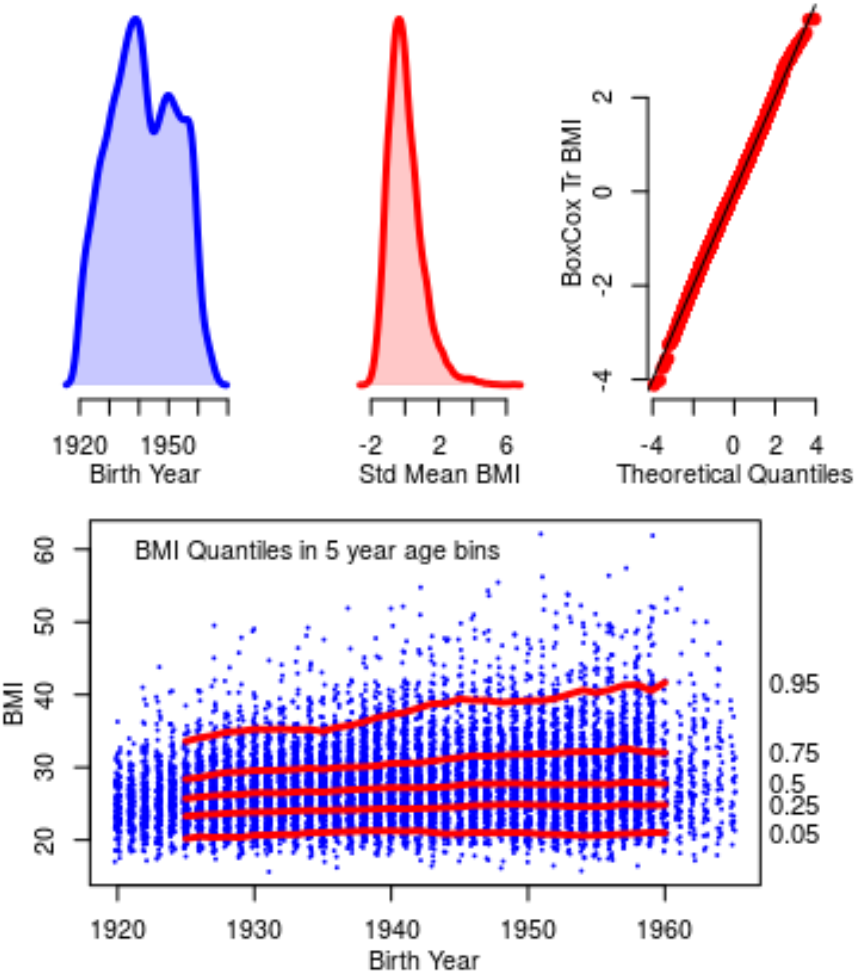
Top: Density of BMI and birthyear in HRS data as well as QQ plot of the Box-Cox transformed version (meant to reduce skew). We standardize respondent’s average BMI taken over all waves; this mean BMI is skewed (skew equals 1.14). This is a common finding with respect to BMI (see, for example, the Figure in [9]). Thus, we also consider a version of this variable in which it has been transformed, via the Box-Cox transformation, to closely approximate a normal distribution. All variables (birthyear, polygenic score, and outcome) are standardized (to have zero mean and unit variance) prior to analysis. Bottom: Illustration of increasing variance in BMI distribution as a function of birth year for analytic sample.

#### E.2 UKB data

We used data from UK Biobank to conduct a series of SNP-level gene-by-environment association analyses. Genotyping and imputation procedures were performed by the original UK Biobank investigators, as described in a previous publication [10].^3^ In the present study, we applied additional thresholds to minimize potential confounding, removing participants with (i) low call rate, (ii) extreme heterozygosity, (iii) chromosomal aneuploidy, (iv) discordant self-reported and chromosomal sex, or (v) missing data for the relevant outcome and covariates. We further focused our analyses on unrelated participants of non-Hispanic European ancestry to minimize the threat of population stratification.

Rather than perform a genome-wide scan of gene-by-environment interactions, we focused on our analyses on loci that have been previously associated with BMI. Specifically, we selected the 97 loci identified in a large genome-wide association study of body mass index that did not include UK Biobank [11]. We successfully identified and extracted the lead SNP for 96/97 (99%) of these loci. For imputed SNPs, dosages were converted to hard calls. Any converted hard calls with a certainty less than .90 were coded as missing.

As some UK Biobank participants had multiple observations for BMI, we computed the mean BMI for each individual. We applied a Box-Cox transformation to reduce skew. Box-Cox transformed BMI was then adjusted for sex, batch, and the first 40 principal components of ancestry, which we estimated ourselves [12]. All variables (SNP, birth year, and body mass index) were standardized to have mean zero and unit variance prior to analysis. The final analytic samples size was N=380,605.

### F Supplemental Resultse

#### F.1 Polygenic score analysis

To better illustrate the differences in the sex-stratified analyses, we consider a graphical analysis inspired by the notion of posterior predictive checks [13]. Using estimates from the empirical data, we construct simulated data sets using Eqn 13 and Eqn 1 and the HRS data. We then consider the predicted percentage of phenotypic variance explained by the genetic predictor across birthyears. Based on Eqn 8, we compute this percentage as

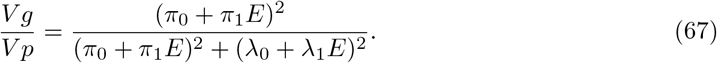

Results of this graphical analysis are in Figure 2. For males (top row), the empirical data (in red) produce a pattern that falls well within the distribution of results from data simulated from the scaling model, but departs more appreciably from results from data simulated under the conventional homoscedastic GxE model. The proportional contribution of the PGS to BMI does not significantly increase with birth year for males. For females, there is more change in penetrance than would be anticipated under the scaling model but less change than would be anticipated under the conventional homoscedastic GxE model. As compared to males,the proportional contribution of the PGS to BMI significantly increases with birth year for females. Thus, for females, the full heteroscedasticity model, in which the contributions of PGS and non-PGS factors independently shift with birth year, may be the most appropriate analytic tool.

We also examined the potential for genetic heteroscedasticity using the genetic heteroscedasticity model (Eqn 44). We observe evidence for moderation of residual variance in BMI by the BMI PGS, as indicated by the λ_2_ parameter. Interestingly, this result holds even for the Box-Cox transformed version of BMI, suggesting that the heteroscedasticity is not entirely an artifact of mean-variance associations induced by the skewed distribution of BMI. One interpretation of these results is that the polygenic score for BMI also captures some degree of “plasticity” (in the sense of [14]). Results of the *ξ_G_* tests (see final column of Table 2) are similar to those of the *ξ_E_* in that we cannot reject the null of the scaling model for the male respondents.

**Table 2:** Estimates from parameters of the full heteroscedasticity model (Eqn 49) in analysis of GxE for BMI as a function of birth year in the HRS. The *ξ_E_* and *ξ_G_* estimates reported are obtained from the environmental and genetic heteroscedasticity models, respectively. We show probabilities for parameters when the maximal probability in a column is larger than 1*e* – 6.

| Gender | BMI<sup>a</sup> | N | $\tau_1$ | $\pi_0$ | $\pi_1$ | $\text{Pr}_{\pi_1}$ | $\lambda_0$ | $\lambda_1$ | $\lambda_2$ | $\text{Pr}_{\lambda_2}$ | $\text{Pr}_{\xi_E}$ | $\text{Pr}_{\xi_G}$ |
| --- | --- | --- | --- | --- | --- | --- | --- | --- | --- | --- | --- | --- |
| All | Std | 11586 | 0.19 | 0.25 | 0.06 | 1.78e-11 | 0.93 | 0.12 | 0.09 | 7.44e-50 | 1.76e-02 | 7.89e-04 |
| All | Trans | 11586 | 0.17 | 0.26 | 0.04 | 1.79e-05 | 0.95 | 0.06 | 0.02 | 1.14e-03 | 2.68e-02 | 5.15e-04 |
| M | Std | 5022 | 0.19 | 0.26 | 0.04 | 4.69e-04 | 0.93 | 0.12 | 0.09 | 1.63e-22 | 3.91e-01 | 2.75e-01 |
| M | Trans | 5022 | 0.17 | 0.26 | 0.03 | 2.11e-02 | 0.95 | 0.07 | 0.03 | 1.33e-03 | 3.83e-01 | 1.65e-01 |
| F | Std | 6564 | 0.20 | 0.25 | 0.06 | 6.65e-09 | 0.93 | 0.12 | 0.09 | 3.52e-31 | 2.24e-02 | 1.21e-03 |
| F | Trans | 6564 | 0.18 | 0.26 | 0.04 | 2.08e-04 | 0.94 | 0.06 | 0.02 | 1.11e-02 | 3.82e-02 | 1.75e-03 |

<sup>a</sup> Std denotes the standardized mean BMI shown in the middle panel of Figure E.1. Trans denotes the Box-Cox transformation.

#### F.2 SNP analysis

Tables F.1 and F.2 contain results discussed in main text.

**Table F.1:** vQTL discoveries for BMI. From the 96 marker SNPs for genome-wide significant loci for BMI in [11], those reaching significance at *p* < .05/96 for moderation of variance, as indicated by the λ_2_ coefficient from the full heterscedasticity model. Parameters are reported for the full heteroscedasticity model, with *ξ_E_* and *ξ_G_* parameters from the environmental and genetic heteroscedasticity models, respectively

| SNP | $\pi_0$ | $\pi_1$ | $\text{Pr}_{\pi_1}$ | $\lambda_0$ | $\lambda_1$ | $\lambda_2$ | $\text{Pr}_{\lambda_2}$ | $\text{Pr}_{\xi_E}$ | $\text{Pr}_{\xi_G}$ |
| --- | --- | --- | --- | --- | --- | --- | --- | --- | --- |
| rs2207139_A | -0.02 | -0.00 | 3.76e-03 | 0.99 | 0.04 | -0.00 | 8.93e-05 | 1.39e-02 | 3.84e-03 |
| rs7903146_C | 0.01 | -0.00 | 6.48e-01 | 0.99 | 0.04 | 0.01 | 3.22e-06 | 8.09e-01 | 9.02e-01 |
| rs9925964_A | 0.02 | -0.00 | 9.07e-01 | 0.99 | 0.04 | 0.00 | 1.10e-04 | 6.11e-01 | 7.50e-01 |
| rs1558902_T | -0.05 | -0.01 | 2.12e-06 | 0.99 | 0.04 | -0.01 | 1.76e-17 | 2.96e-04 | 5.14e-07 |
| rs6567160_T | -0.03 | -0.00 | 1.65e-02 | 0.99 | 0.04 | -0.01 | 2.84e-08 | 8.97e-02 | 1.40e-02 |
| rs2287019_C | 0.02 | -0.00 | 8.71e-01 | 0.99 | 0.04 | 0.01 | 8.15e-06 | 8.27e-01 | 6.66e-01 |

**Table F.2:** Replication of main effects at *p* < .05/96 for the 96 marker SNPs for the genome-wide significant loci for BMI in [11]. Main effects (*π*_0_) come from the full heteroscedasity model.

| SNP | $\pi_0$ | SE($\pi_0$) | Pr$_{\pi_0}$ | Replication? |
| --- | --- | --- | --- | --- |
| rs977747_G | -0.0119 | 0.0016 | 1.3380e-13 | TRUE |
| rs657452_G | -0.0116 | 0.0016 | 6.3858e-13 | TRUE |
| rs11583200_T | -0.0103 | 0.0016 | 1.8037e-10 | TRUE |
| rs3101336_C | 0.0152 | 0.0016 | 1.8616e-21 | TRUE |
| rs12566985_A | -0.0133 | 0.0016 | 1.3812e-16 | TRUE |
| rs12401738_G | -0.0103 | 0.0016 | 1.4297e-10 | TRUE |
| rs11165643_T | 0.0125 | 0.0016 | 6.5757e-15 | TRUE |
| rs17024393_T | -0.0146 | 0.0016 | 1.8806e-19 | TRUE |
| rs543874_A | -0.0271 | 0.0016 | 2.2391e-63 | TRUE |
| rs2820292_C | 0.0131 | 0.0016 | 2.6984e-16 | TRUE |
| rs13021737_G | 0.0279 | 0.0016 | 1.6738e-68 | TRUE |
| rs10182181_A | -0.0243 | 0.0016 | 2.0290e-51 | TRUE |
| rs11126666_G | -0.0021 | 0.0016 | 1.9697e-01 | FALSE |
| rs1016287_C | -0.0128 | 0.0016 | 1.4586e-15 | TRUE |
| rs11688816_G | 0.0083 | 0.0016 | 3.0389e-07 | TRUE |
| rs2121279_C | -0.0050 | 0.0016 | 2.0019e-03 | FALSE |
| rs1460676_T | -0.0083 | 0.0016 | 2.7992e-07 | TRUE |
| rs1528435_T | 0.0101 | 0.0016 | 2.8226e-10 | TRUE |
| rs17203016_A | -0.0071 | 0.0016 | 1.0494e-05 | TRUE |
| rs7599312_G | 0.0116 | 0.0016 | 9.6039e-13 | TRUE |
| rs492400_T | -0.0072 | 0.0016 | 7.8542e-06 | TRUE |
| rs2176040_G | -0.0014 | 0.0016 | 3.7490e-01 | FALSE |
| rs6804842_G | 0.0088 | 0.0016 | 5.6154e-08 | TRUE |
| rs2365389_C | 0.0092 | 0.0016 | 1.0286e-08 | TRUE |
| rs3849570_C | -0.0075 | 0.0016 | 2.5934e-06 | TRUE |
| rs13078960_T | -0.0109 | 0.0016 | 1.3418e-11 | TRUE |
| rs16851483_G | -0.0124 | 0.0016 | 1.3755e-14 | TRUE |
| rs1516725_C | 0.0162 | 0.0016 | 4.0089e-24 | TRUE |
| rs10938397_A | -0.0208 | 0.0016 | 1.5818e-38 | TRUE |
| rs17001654_C | -0.0069 | 0.0016 | 2.3970e-05 | TRUE |
| rs13107325_C | -0.0187 | 0.0016 | 2.1694e-31 | TRUE |
| rs11727676_T | 0.0014 | 0.0016 | 3.9716e-01 | FALSE |
| rs2112347_T | 0.0189 | 0.0016 | 5.4850e-32 | TRUE |
| rs7715256_T | -0.0106 | 0.0016 | 3.8649e-11 | TRUE |
| rs205262_A | -0.0185 | 0.0016 | 8.4528e-31 | TRUE |
| rs2033529_A | -0.0132 | 0.0016 | 2.5074e-16 | TRUE |
| rs2207139_A | -0.0209 | 0.0016 | 3.0425e-38 | TRUE |
| rs9400239_C | 0.0098 | 0.0016 | 1.0575e-09 | TRUE |
| rs9374842_T | 0.0059 | 0.0016 | 2.1120e-04 | TRUE |
| rs13201877_A | -0.0048 | 0.0016 | 2.8052e-03 | FALSE |
| rs13191362_A | 0.0086 | 0.0016 | 8.0189e-08 | TRUE |
| rs1167827_G | 0.0155 | 0.0016 | 4.3736e-22 | TRUE |
| rs2245368_T | -0.0133 | 0.0016 | 2.0238e-16 | TRUE |
| rs9641123_G | -0.0052 | 0.0016 | 1.1563e-03 | FALSE |
| rs6465468_G | -0.0032 | 0.0016 | 4.6521e-02 | FALSE |
| rs17405819_T | 0.0137 | 0.0016 | 1.2564e-17 | TRUE |
| rs16907751_C | 0.0074 | 0.0016 | 4.3644e-06 | TRUE |
| rs2033732_C | 0.0065 | 0.0016 | 4.8454e-05 | TRUE |
| rs4740619_T | 0.0140 | 0.0016 | 2.4629e-18 | TRUE |
| rs10968576_A | -0.0149 | 0.0016 | 1.5486e-20 | TRUE |
| rs6477694_T | -0.0088 | 0.0016 | 4.7682e-08 | TRUE |
| rs1928295_T | 0.0083 | 0.0016 | 1.9969e-07 | TRUE |
| rs10733682_G | -0.0101 | 0.0017 | 1.0969e-09 | TRUE |
| rs7899106_A | -0.0086 | 0.0016 | 1.0639e-07 | TRUE |
| rs17094222_T | -0.0070 | 0.0016 | 1.2704e-05 | TRUE |
| rs11191560_T | -0.0085 | 0.0016 | 1.2063e-07 | TRUE |
| rs7903146_C | 0.0094 | 0.0016 | 4.1177e-09 | TRUE |
| rs4256980_G | 0.0113 | 0.0016 | 1.8886e-12 | TRUE |
| rs11030104_A | 0.0219 | 0.0016 | 1.6990e-42 | TRUE |
| rs2176598_C | -0.0136 | 0.0016 | 1.9700e-17 | TRUE |
| rs3817334_C | -0.0179 | 0.0016 | 6.0633e-29 | TRUE |
| rs12286929_G | 0.0113 | 0.0016 | 2.4293e-12 | TRUE |
| rs7138803_G | -0.0179 | 0.0016 | 1.0766e-28 | TRUE |
| rs11057405_G | 0.0122 | 0.0016 | 2.3471e-14 | TRUE |
| rs12429545_G | -0.0125 | 0.0016 | 1.3652e-14 | TRUE |
| rs9540493_G | -0.0085 | 0.0016 | 1.2686e-07 | TRUE |
| rs1441264_A | 0.0126 | 0.0017 | 4.0919e-14 | TRUE |
| rs10132280_C | 0.0143 | 0.0016 | 1.3063e-18 | TRUE |
| rs12885454_C | 0.0110 | 0.0016 | 5.7772e-12 | TRUE |
| rs11847697_C | -0.0072 | 0.0016 | 7.7682e-06 | TRUE |
| rs7141420_T | 0.0145 | 0.0016 | 5.3645e-19 | TRUE |
| rs3736485_G | -0.0082 | 0.0016 | 3.5057e-07 | TRUE |
| rs16951275_T | 0.0159 | 0.0016 | 3.9682e-23 | TRUE |
| rs7164727_T | 0.0104 | 0.0016 | 9.0124e-11 | TRUE |
| rs758747_C | -0.0088 | 0.0016 | 7.5808e-08 | TRUE |
| rs12446632_G | 0.0149 | 0.0016 | 2.1385e-20 | TRUE |
| rs2650492_G | -0.0124 | 0.0016 | 1.7199e-14 | TRUE |
| rs3888190_C | -0.0191 | 0.0016 | 1.1586e-32 | TRUE |
| rs4787491_G | 0.0126 | 0.0016 | 4.9604e-15 | TRUE |
| rs9925964_A | 0.0165 | 0.0016 | 1.0272e-24 | TRUE |
| rs2080454_A | -0.0057 | 0.0016 | 4.8633e-04 | TRUE |
| rs1558902_T | -0.0503 | 0.0016 | 1.0423e-214 | TRUE |
| rs9914578_C | -0.0064 | 0.0016 | 6.7263e-05 | TRUE |
| rs1000940_A | -0.0105 | 0.0016 | 5.9037e-11 | TRUE |
| rs12940622_G | 0.0129 | 0.0016 | 1.1756e-15 | TRUE |
| rs1808579_C | 0.0131 | 0.0016 | 3.3014e-16 | TRUE |
| rs7239883_A | -0.0073 | 0.0016 | 6.8097e-06 | TRUE |
| rs7243357_T | 0.0092 | 0.0016 | 9.8026e-09 | TRUE |
| rs6567160_T | -0.0311 | 0.0016 | 5.0802e-83 | TRUE |
| rs17724992_A | 0.0111 | 0.0016 | 5.1966e-12 | TRUE |
| rs29941_G | 0.0093 | 0.0016 | 7.1063e-09 | TRUE |
| rs2075650_A | 0.0113 | 0.0016 | 1.7586e-12 | TRUE |
| rs2287019_C | 0.0183 | 0.0016 | 4.9748e-30 | TRUE |
| rs3810291_A | 0.0191 | 0.0016 | 8.5285e-33 | TRUE |
| rs6091540_C | 0.0134 | 0.0016 | 1.1496e-16 | TRUE |
| rs2836754_C | 0.0076 | 0.0016 | 1.8681e-06 | TRUE |

## Footnotes

1 We use Pr_*β*_ to refer to probability associated with a test of the null hypothesis that some parameter *β* equals zero.

2 https://github.com/ben-domingue/scalingGxE

3 See also https://www.ukbiobank.ac.uk/scientists-3/genetic-data/.

